# Distributed cortical regions for the recall of people, places and objects

**DOI:** 10.1101/2022.08.03.502612

**Authors:** Alexis Kidder, Edward H Silson, Matthias Nau, Chris I Baker

## Abstract

Human medial parietal cortex (MPC) is recruited during multiple cognitive processes. Previously, we demonstrated regions specific to recall of people or places and proposed that the functional organization of MPC mirrors the category selectivity defining the medial-lateral axis of ventral temporal cortex (VTC). However, prior work considered recall of people and places only and VTC also shows object-selectivity sandwiched between face- and scene-selective regions. Here, we tested a strong prediction of our proposal: like VTC, MPC should show a region specifically recruited during object recall, and its relative cortical position should mirror the one of VTC. While responses during people and place recall showed a striking replication of prior findings, we did not observe any evidence for object-recall effects within MPC, which differentiates it from the spatial organization in VTC. Importantly, beyond MPC, robust recall-effects were observed for people, places, and objects on the lateral surface of the brain. Place-recall effects were present in the angular gyrus, frontal eye fields and peripheral portions of early visual cortex, whereas people-recall selectively drove response in the right posterior superior temporal sulcus. Object-recall effects were largely restricted to a region posterior to left somatosensory cortex, in the vicinity of the supramarginal gyrus. Taken together, these data demonstrate that while there are distributed regions active during recall of people, places and objects, the functional organization of MPC does not mirror the medial-lateral axis of VTC but reflects only the most salient features of that axis - namely representations of people and places.

**Significance statement:** Human medial parietal cortex (MPC) is recruited during multiple cognitive processes. Recently, we proposed a framework for interpreting the functional organization of MPC by suggesting that it reflects the categorical preferences for people and places that is evident also in ventral temporal cortex (VTC). Because VTC also exhibits selectivity for objects, we here extend this framework to test whether MPC also shows object selectivity during recall. Robust people and place recall effects were evident in MPC, but we found no evidence for object-recall within MPC, suggesting that MPC and VTC are not mirror-copies of each other. Together, these data suggest that the functional organization of MPC reflects the most salient categorical representations within VTC for people and places.

## Introduction

Human medial parietal cortex (MPC) is associated with a broad array of cognitive functions, including memory recall (Wagner et al., 2005; Vilberg & Rugg, 2008; Kim, 2013; Gilmore et al., 2015; Silson, Gilmore, et al., 2019; Silson, Steel, et al., 2019; Steel et al., 2021) visual scene perception (Baldassano et al., 2013; Epstein & Baker, 2019; Silson, Gilmore, et al., 2019), scene construction (Hassabis et al., 2007), processing of spatial and other contextual associations (Bar & Aminoff, 2003), navigation (Epstein, 2008; Epstein & Baker, 2019), future thinking (Szpunar et al., 2007; Benoit & Schacter, 2015; Gilmore et al., 2018), and mental orientation (Peer et al., 2015).

The diversity with which MPC is recruited has prompted recent work from our group (Silson, Steel, et al., 2019; Steel et al., 2021; Bainbridge et al., 2021) and others (Peer et al., 2015; Chrastil, 2018; Woolnough et al., 2020; Deen & Freiwald, 2021) to explore whether there is an underlying functional organization to MPC. Specifically, in a series of experiments, we identified a functional link between the medial-lateral axis of ventral-temporal cortex (VTC) and the posterior/ventral - anterior/dorsal axis of MPC (Silson, Steel, et al., 2019). First, distinct regions of MPC were identified based upon patterns of differential connectivity with face- and scene-selective regions, respectively. Second, we found that these same regions responded most strongly when processing visual stimuli of their preferred category. Finally, via cued recall of either people or places, we demonstrated an alternating pattern of recruitment along the posterior/ventral - anterior/dorsal axis of MPC. These data led to the suggestion that the functional organization of MPC may be a recapitulation of the medial-lateral axis of VTC **(Figure 1A)**, in which visual face selectivity (i.e. fusiform face area, FFA) and visual scene selectivity (i.e. parahippocampal place area, PPA) **(Figure 1B)** (Kanwisher & Dilks, 2013; Weiner et al., 2014; Epstein & Baker, 2019) are located in distinct, yet adjacent regions. Importantly VTC also contains a region that has been characterized as object-selective (i.e. posterior fusiform cortex, pFS), in-between FFA and PPA. Here, we tested a strong prediction of our framework and investigated whether there is also an object-selective region in MPC during memory recall.

**Figure 1:**
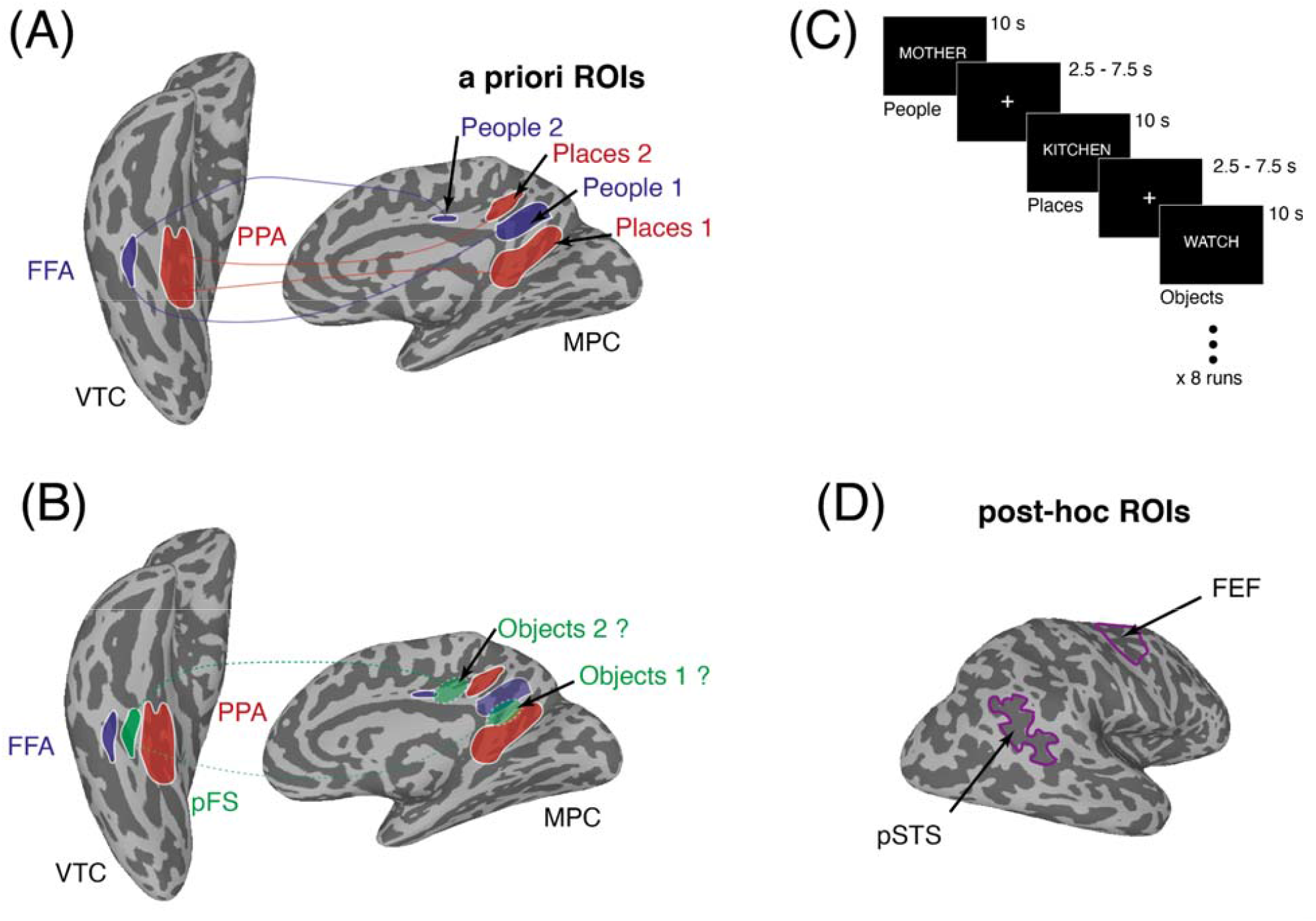
Functional link between VTC and MPC, predictions, task schematic & post-hoc ROIs. **(A)** Partially inflated ventral and medial views of the right hemisphere of a representative participant are shown (light gray = gyri, dark gray = sulci). Overlaid in false colour onto the ventral surface are group-based ROIs for face-selective FFA (Blue) and scene-selective PPA (Red). Overlaid in false colour onto the medial surface are group-based ROIs for people and place recall areas within MPC (People 1 / People 2 = Blue; Places 1 / Places 2 = Red) taken from our prior work (Silson, Steel, et al., 2019). Along the posterior/ventral - anterior/dorsal axis of MPC these regions show an alternating pattern (Places, People, Places, People). Lines connecting VTC with MPC illustrate the consistent category preferences. **(B)** The same ventral and medial views of the right hemisphere in (A) are shown. Overlaid in false colour onto the ventral surface is the group-based ROI for object-selective pFs (Green). Within VTC, object-selective responses (pFS) are sandwiched between face-selective responses (FFA(more laterally and scene-selective responses (PPA) more medially. If the functional organization of MPC reflects that of VTC, we predict object-recall areas to be located in-between areas recruited during people and place recall. This predicted pattern of results in overload in false colour onto the medial surface. Green dashed lines depict the hypothesized link with pFs on the ventral surface. **(C)** Task schematic. During each trial participants were instructed to recall from memory either personally familiar people (e.g. Mother), places (e.g. Kitchen) or objects (e.g. Watch). Participants were instructed to visualise the target as vividly as possible for the duration of the trials (10 s). Trials were separated by a variable inter-trial-interval (2.5-7.5s). Each run contained six randomised trials from each category. Participants completed 8 runs of the experiment. **(D)** Overlaid onto a lateral view of the right hemisphere are the locations of two post-hoc ROIs (purple outlines): the pSTS was taken from an independent group-based localizer (see methods). Frontal eye fields were defined according to a probabilistic retinotopic atlas (Wang et al., 2015).

Participants recalled personally familiar people, places or objects, allowing us to accomplish two goals. First, it provided an opportunity for a direct, yet independent test of prior work on recall of people and places in MPC (Silson, Steel, et al., 2019). Second, it allowed us to examine whether or not the recall of personally familiar objects recruited distinct regions of MPC, akin to the perception of objects in VTC (i.e. pFS). Given the stereotypical location of pFS relative to both FFA and PPA in VTC (Kanwisher & Dilks, 2013), we hypothesized that if the recall of objects recruited distinct regions of MPC these would likely fall between those recruited during place and people recall (Silson, Steel, et al., 2019) **(Figure 1B)**.

Recall of people and places produced a pattern of recruitment within MPC that was largely indistinguishable from prior work (Silson, Steel, et al., 2019) and constitutes a complete yet independent replication. However, we found no evidence for distinct responses during object-recall within MPC, despite strong predictions based on the functional organization of VTC. However, we did observe object-recall effects outside MPC, in a region posterior and inferior of primary somatosensory cortex in the vicinity of the supramarginal gyrus. Moreover, people recall selectively drove responses in the right-hemispheric posterior superior temporal sulcus (pSTS), whereas place recall recruited bilateral angular gyrus, peripheral portions of early visual cortex (EVC) and frontal eye fields. Taken together these data highlighted distributed regions active during recall and suggest that the functional organization of MPC is largely dominated by the representation of people and places with object-recall if anything engaging action-related areas, which contrasts it to VTC despite clear parallels.

## Methods

### Participants

Twenty-four participants (17 female, mean age = 24.2 years) were recruited from the DC area and NIH community. All participants were right-handed with normal or corrected-to-normal vision and neurologically healthy. All participants gave written informed consent according to procedures approved by the NIH Institutional Review Board (protocol 93 M-0170, clinical trials # NCT00001360). Participants were compensated monetarily for their time. The sample size for the memory experiment was based on prior work from our group employing a very similar paradigm (Silson, Gilmore, et al., 2019; Silson, Steel, et al., 2019), where sample sizes were n=19 and n=24, respectively. Due to excessive motion during scanning only 20 out of the 24 datasets were analyzed in full.

### Stimuli and tasks

#### Memory Experiment

Experimental details were based on prior work (Silson, Steel, et al., 2019) and stimuli consisted of written names of 12 personally familiar people (e.g. Mother), 12 personally familiar places (e.g. Kitchen) and 12 personally familiar objects (e.g. Watch). The stimuli were provided by participants through a survey completed prior to the fMRI scan. Importantly, we selected objects that did not have strong contextual associations to certain places/scenes (e.g. wallet), as contextual association is thought to drive responses in certain MPC regions (Bar & Aminoff, 2003). Word stimuli were presented in white 18-point Arial font, all capital type against a black background. During each trial of the task **(Figure 1C)**, participants were instructed to visualize the named item from memory as vividly as possible for the duration of the trial (10 s). Mean character length: Objects = 6.31, Places = 12.05, People = 6.42. Trials were separated by a variable inter-trial interval (2.5–7.5 s). In each of the eight runs, there were six trials of each condition (People, Places, Objects) presented in a randomized order, for a total of 18 trials per run (144 trials total). Across the entire scanning session each stimulus was presented four times.

#### Post scan questionnaire

After the scan, participants completed a questionnaire in which they rated how vividly they were able to visualize from memory each item named during the memory runs. The stimuli were listed in the same order they appeared during the Memory Experiment and were rated on a 4-point Likert type scale (1 = not at all vivid; 4 = extremely vivid). If the participant could not visualize the stimulus at all while in the scanner, they checked a box on the questionnaire.

##### Functional imaging parameters

###### Memory experiment

All scans were performed on a 3.0T GE 750 MRI scanner using a 32-channel head coil. All functional images were acquired using a BOLD-contrast sensitive three-echo echo-planar sequence (ASSET acceleration factor = 2, TEs = 12.5, 27.7, and 42.9 ms, flip angle = 75°, 64 × 64 matrix, in-plane resolution = 3.2×3.2 mm, slice thickness = 3.5 mm, TR=2500ms, 35 slices).

### fMRI data preprocessing

Data were preprocessed using AFNI (Cox, 1996) (RRID: SCR_005927). The first 4 volumes of each run were discarded to allow for T1 equilibration effects. Initial preprocessing steps for fMRI data were carried out on each echo separately. Slice-time correction was applied (3dTShift) and signal outliers were attenuated (3dDespike). Motion-correction parameters were estimated from the middle echo based on rigid-body registration of each volume to the first volume of the scan; these alignment parameters were then applied to all echos. Data from all three acquired echoes were then registered to each participants’ T1 image and combined to remove additional thermal and physiological noise using multi-echo independent components analysis (ME-ICA, Kundu et al., 2012, 2013). This procedure computes a weighted-average of the three echo times for each scan run to reduce thermal noise within each voxel. It subsequently performs a spatial ICA to identify and remove noise components from the data. This is accomplished by comparing each component to a model that assumes a temporal dependence in signal decay (i.e., that is ‘BOLD-like’) and to a different model that assumes temporal independence (i.e., that is ‘non-BOLD-like’). Components with a strong fit to the former and a poor fit to the latter are retained for subsequent analysis. This procedure was conducted using default options in AFNI’s tedana.py function. ME-ICA processed data from each scan were then aligned across runs for each participant.

### Memory analysis

Analyses were conducted using a general linear model (GLM) and the AFNI programs 3dDeconvolve and 3dREMLfit. The data at each time point were treated as the sum of all effects thought to be present, and the time series was compared against a Generalized Least Squared (GLSQ) regression model fit with REML estimation of the temporal auto-correlation structure. Responses were modeled by convolving a standard gamma function with a 10 s square wave for each condition of interest (People, Places, Objects). Estimated motion parameters were included as additional regressors of non-interest, and fourth-order polynomials were included to account for slow drifts in the MR signal over time. Significance was determined by comparing the beta estimates for each condition (normalized by the grand mean of each voxel for each run) against baseline (fixation only).

### Sampling of data to the cortical surface

In each participant, statistical datasets were projected onto surface reconstructions of each individual participant’s hemispheres using the Surface Mapping with AFNI (SUMA) software (Saad & Reynolds, 2012). First, data were aligned to high-resolution anatomical scans (align_epi_anat.py). Once aligned, these data were projected onto the cortical surface (3dVol2Surf) and smoothed with a 2 mm full-width-at-half-maximum 2D Gaussian kernel.

### Cortical Regions of Interest

Initially, we utilized the whole brain data from prior work (Silson, Steel, et al., 2019) to define four regions of interest (ROIs) within MPC of each hemisphere (Places1, People1, Places2 & People2, see **Figure 1A**). Importantly, these data were acquired in an independent group of participants. The frontal-eye-fields (FEF) were defined using a probabilistic retinotopic atlas (Wang et al., 2015). In addition, we made use of a group-based (n=15) dynamic localizer using the contrast of faces > objects from an unpublished dataset in independent participants to localize face-selective pSTS.

### Eyeball-voxel analysis

To test whether recall-related eye movements may have contributed to our imaging results, we performed multivariate pattern analysis (MVPA) using the multi-voxel pattern of the eyeballs. To do so, we identified the eyeball voxels in our images following established approaches (Frey & Nau et al., 2021), which entailed co-registering the eyeballs to the ones of a template in which the eyes were manually delineated. This dedicated eyeball coregistration procedure ensured that the eyeball voxels were correctly identified and that the corresponding multi-voxel pattern could be extracted from each functional volume (see (Frey & Nau et al., 2021) for more details). To reduce noise, we then removed voxels with low temporal signal-to-noise ratio, excluding the lower 50 percent, and we linearly detrended the time series of each voxel followed by Z-scoring. We then median-averaged all functional volumes corresponding to the same trial, obtaining one multi-voxel-pattern per trial, which served as the basis for MVPA. Specifically, we computed pairwise Spearman correlations between all trials of a given category (e.g. places vs. places) as well as across categories (e.g. places vs. objects and people). These comparisons were cross-validated across runs to avoid effects of autocorrelated time-series noise (i.e. we excluded comparisons of trials that were tested in the same run). If eye movements were more similar between trials of the same category than between trials of different categories, this should result in overall higher Spearman correlations for the within-category comparisons compared to the across-category comparisons. We tested this possibility using one-tailed paired t-tests. In addition, because averaging across volumes may reduce sensitivity to eye-movement effects within each trial, we further repeated these analyses for each of the four individual volumes acquired in each trial.

### Linear mixed effects analysis

To look at the whole brain memory effects we employed a linear-mixed-effects model (3dLME) in each hemisphere separately. The model comprised a single factor: Category (People, Places, Objects).

### Statistical approach

ROI statistical analyses were performed using R Studio software package (version 1.1). For all analyses we conducted repeated measures analysis of variance (rmANOVA). When Mauchley’s test of sphericity was violated, main effects and interactions were corrected using the Greenhouse-Geisser (GG) correction to allow appropriate interpretation. When a significant three-way interaction was observed, we performed two separate two-way ANOVAs to explore the nature of the interaction. When a significant two-way interaction was observed, post-hoc paired *t*-tests were conducted and corrected for multiple comparisons using Bonferroni correction.

## Results

Our goals here were two-fold. First, we aimed to replicate prior findings of distinct regions in MPC for the recall of people and places. Second, we explored whether any additional region could be identified via selective recruitment during the recall of personally familiar objects. Prior work (Silson, Steel, et al., 2019) highlighted the functional link between the medial-lateral axis of VTC and the ventral-dorsal axis of MPC for perceptual and mnemonic representations of people and places. Here, we extended this work to include recall of personally familiar objects to see if a) recall of such items recruited a spatially distinct area of MPC and b) if so, whether this area was located in-between the place and people regions as predicted based on the functional organization of VTC **(Figure 1A, 1B)**. Below we focus initially on our whole-brain and a priori ROI analyses within MPC itself, before exploring memory recall effects outside of MPC at both the whole-brain level and within our post-hoc ROIs.

### Replication of people and place recall within MPC

**Figure 2A** depicts the whole-brain contrast of Places versus People (p=6.3-5, nodewise q=9.2-4) across medial views of both hemispheres, with MPC enlarged. An alternating pattern of people and place recall was present throughout MPC in both hemispheres. Indeed, the topography of these regions closely matches those reported in prior work despite originating from an independent group of participants. To explore these data further, we calculated the mean response (t-value versus baseline) to each category within a priori ROIs (People 1, Places 1, People 2 & Places 2, **Figure 2B**), which we split into two sets: a larger more ventral pair (i.e., People 1, Places 1) and a smaller more dorsal pair (i.e., People 2, Places 2), based on prior work (Silson, Steel et al., 2019). A consistent pattern was observed within the a priori ROIs and we discuss each pair of regions in turn.

**Figure 2:**
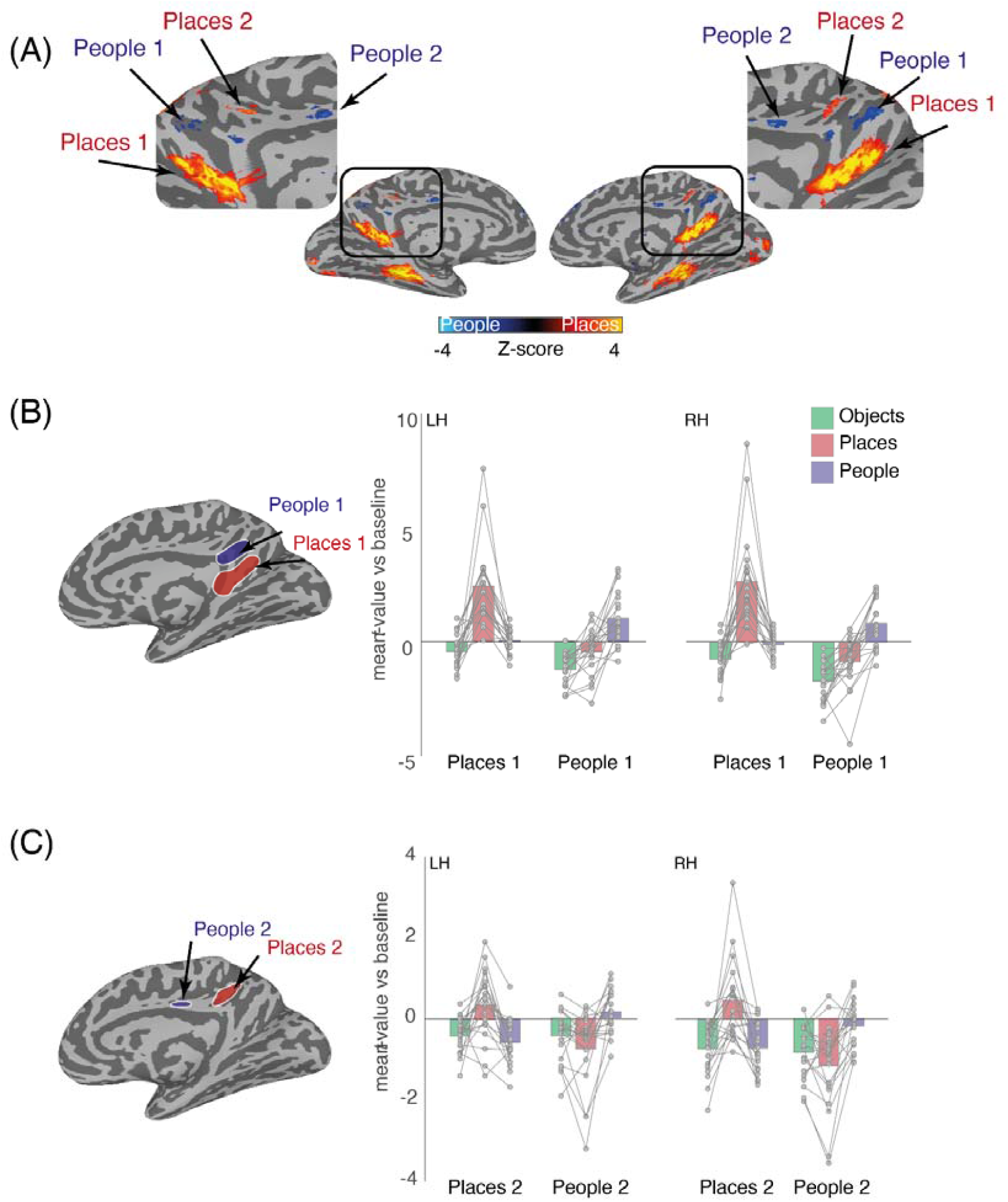
Recall effects in MPC and a priori ROIs. **(A)** Partially inflated medial views of both hemispheres are shown. The contrast of People versus places (p=6.3-5, nodewise q=9.2-4, corrected for multiple comparisons) is overlaid with cold colors representing regions of MPC recruited during People recall and hot colors representing regions of MPC recruited during place recall. Consistent with prior work, an alternating pattern of people and place recall is evident throughout MPC. Enlarged versions of MPC are inset and demonstrate the alternating pattern. **(B)** A partially inflated medial view of the right hemisphere is shown with a priori Places 1 (blue) and People 1 (red) ROIs overlaid. Bars represent the mean group-averaged response to each category (Objects=green, Places=red, People=blue) in both hemispheres. Individual participant data points are plotted and connected. In both hemispheres, recall of places elicited the largest response in Places 1 with responses during people or object recall being close to or below zero. In People 1, responses were maximally positive during people recall, but negative during both object and place recall in both hemispheres. **(C)** Same as (B) but for Places 2 and People 2. Overall, a similar pattern of results is evident, although smaller in magnitude. Place recall maximally drove responses in Places 2 (bilaterally), whereas people recall maximally drove responses in People 2 (bilaterally).

First, recall of places selectively recruited Places 1 and the recall of people selectively recruited People 1 bilaterally **(Figure 2B)**. Interestingly, the recall of objects produced negative responses in both regions, but to varying degrees with responses more negative during object-recall in People 1. To quantify these effects, we conducted a three-way rmANOVA with Hemisphere (Left, Right), ROI (Places1, People1) and Category (Objects, Places, People) as within-participant factors. The main effects of Hemisphere (F(1, 19)=10.21, p=0.004), ROI (F(1, 19)=25.79, p=6.66-5) and Category (F(2, 38)=29.74, p=1.68-8) were all significant, reflecting on average larger responses in the right hemisphere, in Places 1 and for the recall of places, respectively. These were qualified however by significant interactions of Hemisphere x ROI(F(1, 19)=6.16, p=0.02), Hemisphere x Category(F(2, 38)=3.74, p=0.03), ROI x Category(F(2, 38)=42.66, p=3.93-8, GG-corrected) and Hemisphere x ROI x Category(F(2, 38)=5.24, 0.009).

To better interpret the three-way interaction, we performed two two-way rmANOVAs in each hemisphere separately. In the left hemisphere, the main effects of ROI (F(1, 19)=20.21, p=0.0002) and Category (F(2, 38)=29.26, p=2.02-8) were significant, again reflecting larger responses in Places 1 over People 1 and for the recall of places over either people or objects. Importantly, these main effects were qualified by a significant ROI x Category interaction (F(2, 38)=34.38, p=1.37-7, GG-corrected), reflecting the differential pattern of responses across the two ROIs. In the right hemisphere, the main effects of ROI (F(1, 19)=27.95, p=4.19-5) and Category (F(2, 38)=28.07, p=8.59-7, GG-corrected) were significant (larger responses in Places 1 and for places over people or objects), and were qualified by a significant ROI x Category interaction (F(2, 38)=47.46, p=2.36-8, GG-corrected), reflecting the differential pattern of responses across the two ROIs. A series of paired t-tests (Bonferroni corrected, alpha=0.004) confirmed greater responses in Places 1 bilaterally during place recall as compared to either people (LH: t(19)=5.89, p=1.12-5; RH: t(19)=5.84, p=1.25-5) or objects (LH: t(19)=5.99, p=8.99-6); RH: t(19)=6.01, p=8.73-6). The response to people was also different to objects in the right hemisphere (t(19)=3.28, p=0.003), but not the left (t(19)=2.30, p=0.03). In People 1, responses during people recall were greater than both places (LH: (t(19)=4.63, p=0.001; RH: t(19)=6.31, p=4.60-6) and objects (LH: t(19)=6.64, p=2.34-6; RH: t(19)=7.87, p=2.11-7). The response to places was greater than the response to objects in the left hemisphere (t(19)=3.74, p=0.001), but not the right (t(19)=2.99, p=0.007).

Second, we replicated a striking finding of previous work: another more dorsal pair of regions also appear to be selectively recruited during people and place recall (i.e., People 2, Places 2). Here, like previously, selective responses were observed in both ROIs despite lower overall magnitudes relative to their larger, more ventral counterparts. That is, recall of places still maximally drove positive responses in Places 2 whereas recall of people resulted in the most positive responses in People 2. Again, responses during object recall were almost always negative across both ROIs (**Figure 2C)**. Consistent with the analyses described above, we again conducted a 3-way rmANOVA, revealing main effects of Hemisphere (F(1, 19)=10.09, p=0.004) and Category (F(2, 38)=3.37, p=0.04), reflecting on average larger responses in the right hemisphere and in People 2. There was no main effect of ROI (F(1, 19)=3.28, p=0.08). Only the ROI x Category interaction (F(2, 38)=30.94, p=5.29-7, GG-corrected) was significant (p>0.05, in all other cases). Given that hemisphere did not interact with the other factors, we collapsed across hemispheres and ran a two-way rmANOVA. The main effect of ROI (F(1, 19)=3.28, p=0.08) was not significant, but the main effect of Category (F(2, 38)=3.37, p=0.04) was, as was the ROI x Category interaction (F(2, 38)=20.94, p=5.29-7, GG-corrected). Again, a series of paired t-tests (Bonferroni corrected, alpha=0.008) revealed greater responses during place recall in Places 2 than either people (t(19)=5.72, p=1.63-5) or objects (t(19)=4.48, p=0.0002). The response to people was not different to objects (t(19)=0.51, p=0.61). In People 2, people recall was greater than recall of both places (t(19)=4.33, p=0.0003) and objects (t(19)=3.61, p=0.001). The recall of places was not different from objects (t(19)=1.91, p=0.07).

### No evidence for object-recall within MPC

Having replicated prior work on people and place recall, we next examined object recall. Based on the topography of categorical preferences along the medial-lateral axis of VTC **(Figure 1B)**, we predicted that object recall would also recruit distinct region(s) of MPC in between those recruited during people and place recall. However, there was a striking lack of object-related activity within MPC. Indeed, when contrasting object recall directly with either places **(Figure 3A)** or people **(Figure 3B)** no suprathreshold object-recall clusters are evident, whereas clusters recruited during place and people recall are easily identifiable and closely match those highlighted in **Figure 2A**. We next asked whether there are object-recall effects outside of MPC.

**Figure 3:**
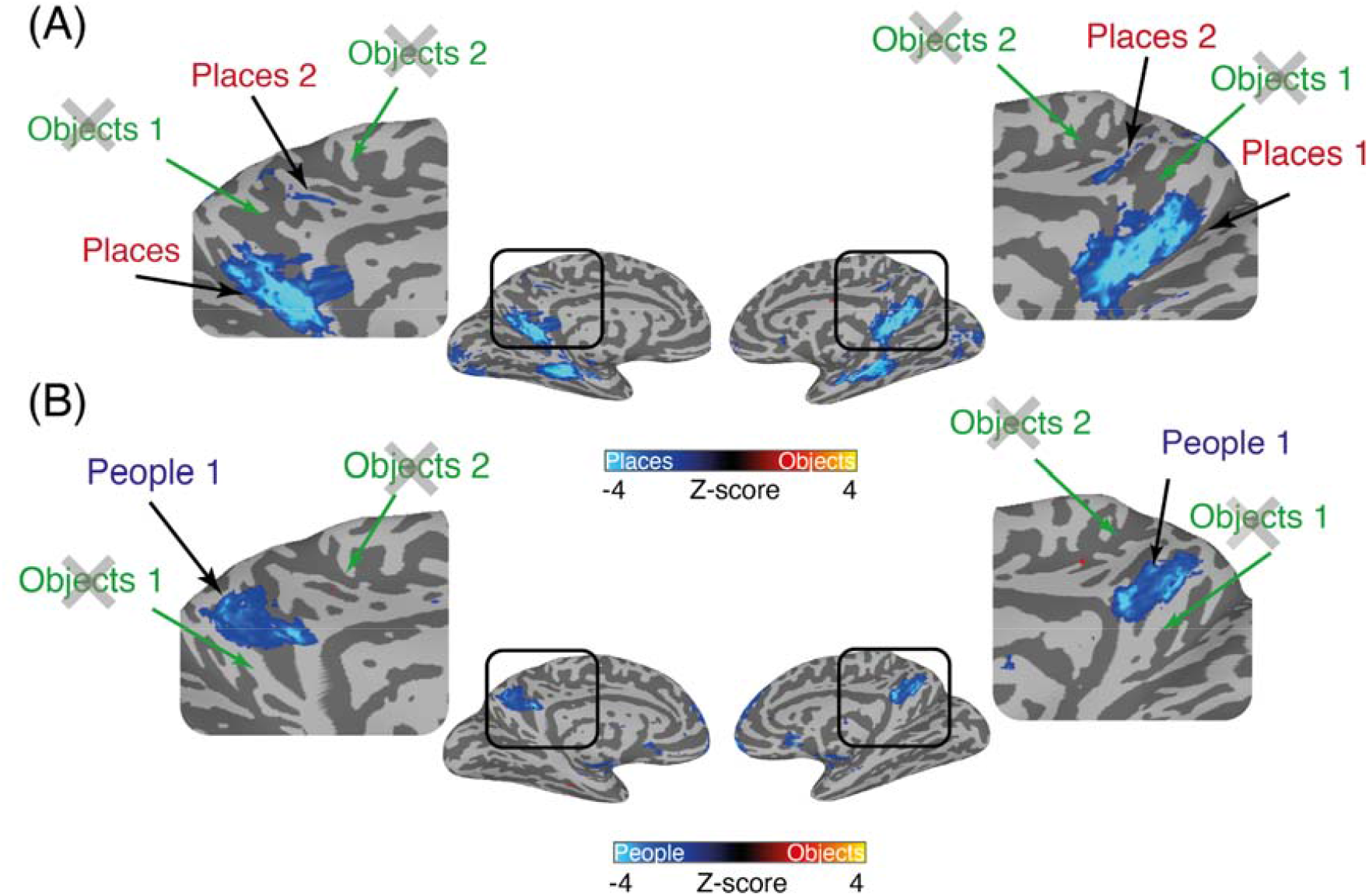
Lack of object recall within MPC. **(A)** Partially inflated medial views of both hemispheres are shown. The contrast of Places versus Objects (p=6.3-5, nodewise q=9.2-4, corrected for multiple comparisons) is overlaid with cold colors representing regions of MPC recruited during Place recall and hot colors representing regions of MPC recruited during Object recall. No suprathreshold object clusters are present in the predicted locations, whereas place recall clusters are easily identifiable. **(B**) Same as but for the contrast of People versus Objects. Again, there is a lack of object-recall but clear people recall.

### Distinct category-recall effects outside of MPC

Prior work (Gilmore et al., 2015; Peer et al., 2015; Silson, Gilmore, et al., 2019; Silson, Steel, et al., 2019; Favila et al., 2019; Steel et al., 2021; Bainbridge et al., 2021) reported that memory-recall effects were not restricted to MPC and could also be found on both the lateral surface of occipitotemporal cortex and in frontal regions. **Figure 4A** depicts the whole-brain effect of Category on lateral views of both hemispheres (p=8.7-5, nodewise q=9.0-4). Significant effects are present near the postcentral sulcus of the left hemisphere, in posterior angular gyrus (bilaterally), the posterior superior temporal sulcus (pSTS) in the right hemisphere, and the dorsal frontal lobe (bilaterally), likely corresponding to the frontal eye fields (FEF). To examine these effects in more detail, we adopted the same approach taken above and next explored the category contrasts. **Figure 4B** depicts the contrast of Places versus People across the same lateral views. Lateral and frontal memory-recall responses are dominated by place recall, in particular in the angular gyrus and the (putative) FEF. A small significant cluster of people-related recall was identified within the pSTS in the right hemisphere only. **Figure 4C-D**, depicts the contrast of Object versus Places and Objects versus People across the same lateral views. Unlike MPC, we did observe significant object-related clusters here, particularly within inferior and posterior portions of the postcentral sulcus in the left hemisphere in the vicinity of the supramarginal gyrus (particularly prominent for the Objects versus People contrast). In the following sections, we explore these effects further in relation to prior studies, in particular sampling the mean response to each category from post-hoc ROIs, corresponding to the face-selective pSTS (see methods), and the FEF (Wang et al., 2015).

**Figure 4:**
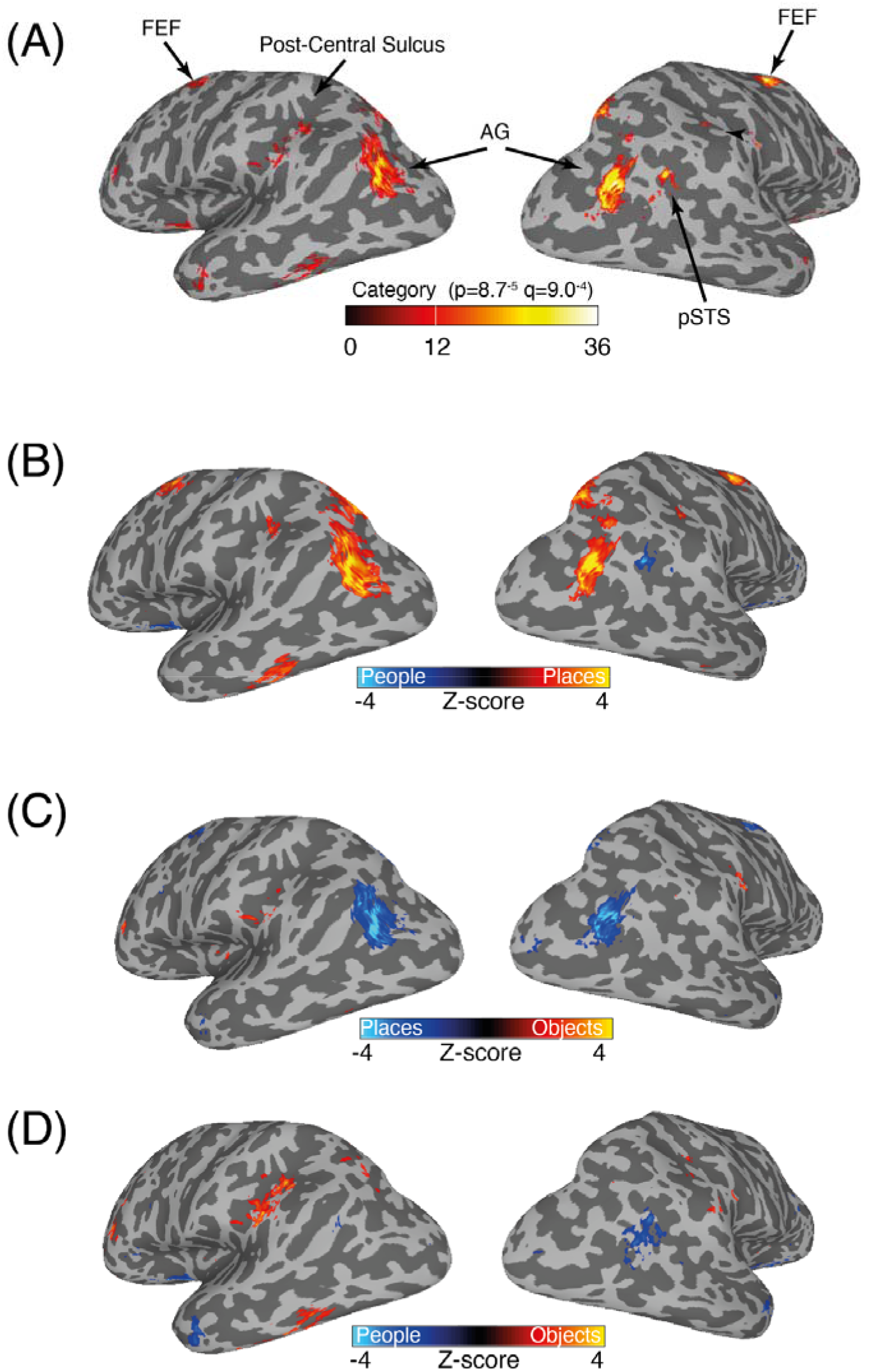
Recall effects outside MPC. **(A)** Partially inflated lateral views of both hemispheres are shown. The main effect of Category (p=8.7-5, nodewise q=9.0-4, corrected for multiple comparisons) is overlaid in false-color. Memory recall effects are present within posterior of the post-central sulcus, within the AG and putative FEF bilaterally and within pSTS of the right hemisphere. **(B)** The contrast of People versus places (p=6.3-5, nodewise q=9.2-4) is overlaid onto the same medial views as (A). Both the AG and FEF were recruited during place recall, whereas people recall recruited pSTS in the right hemisphere. **(C)** Same as (B) but for the contrast of places versus objects. Object-recall effects were evident in regions inferior and posterior to primary somatosensory cortex and in the vicinity of AIP. Place recall effects remained in the AG and FEF. **(D)** Same as (B) but for the contrast of people versus objects. Again, object-recall effects were evident in the SC, particularly in the left hemisphere. People-related recall in right pSTS also remained.

### People recall effects in the posterior STS

A surprising finding of our whole-brain analyses, which we did not observe in our prior study, was a small cluster within the pSTS of the right hemisphere that appeared when we contrasted either people versus places **(see Figure 4B)** or people versus objects **(see Figure 4D)**,. This cluster might correspond to the visual face-selectivity that has previously been reported in pSTS. To better understand the responses we observed, we sampled the mean response to each category from a group-based ROI for face-selective pSTS taken from an independent group of participants (see Methods). Despite responses in this region being negative for all categories **(Figure 5)**, responses were less negative during people recall compared to all other categories. A one-way ANOVA with Category (same levels as above) as the within-participant factor showed a main effect of Category (F(2, 38)=4.22, p=0.02). A series of paired t-tests (Bonferroni corrected, alpha=0.0167) revealed that responses during people recall were different from those during recall of objects (t(19)=2.55, p=0.01), but not places (t(19)=1.40, p=0.19). Responses were not different between places and objects (t(19)=1.72, p=0.10).

**Figure 5:**
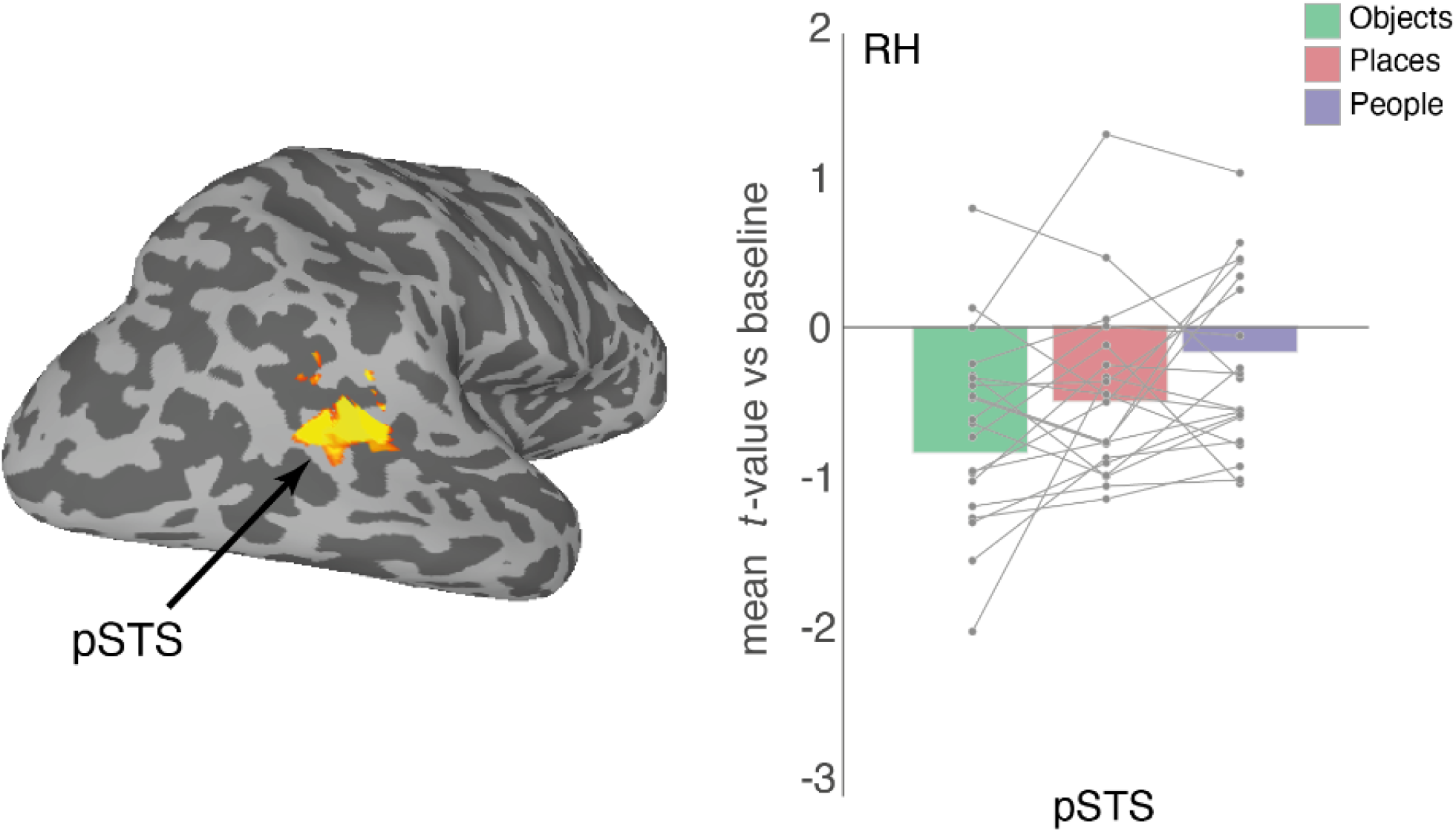
People recall in the right pSTS. A lateral view of the right hemisphere is shown. Outlined in white is the anatomical ROI for right pSTS taken from an independent group-based localizer (see methods). Bars represent the mean response to all categories from the pSTS in the right hemisphere. Individual participant data points are plotted and connected. Despite negative responses to all categories, these responses differentiate in the basis of category with responses during people recall being less negative (relatively more positive)

### Place recall recruits the frontal eye fields

Our whole-brain analyses also highlighted significant category effects in the approximate location of the FEF. In order to quantify this effect and to confirm that it was indeed localized to the FEF, we sampled the mean response to each category from a probabilistic retinotopic FEF mask (Wang et al., 2015) in both hemispheres **(Figure 6)**. A two-way rmANOVA with Hemisphere and Category as factors (same levels as above) revealed main effects of Hemisphere (F(1, 19)=17.41, p=0.0005) and Category (F(2, 28)=11.84, p=0.0001). On average, the magnitude was larger in the left hemisphere, and we observed larger responses during place recall compared to other categories, respectively). The Hemisphere x Category interaction (F(2, 38)=2.09, p=0.13) was not significant. A series of paired t-tests (Bonferroni corrected, alpha=0.008) revealed that responses during place recall were greater in the FEF (bilaterally) than either recall of people (LH: t(19)=3.27, p=0.003; RH: t(19)=4.37, p=0.0003) or objects (LH: t(19)=3.54, p=0.002; RH: t(19)=3.90, p=0.0009). Importantly, responses were not different between people and object recall in either hemisphere (LH: t(19)=0.58, p=0.56; RH: t(19)=1.81, p=0.08).

**Figure 6:**
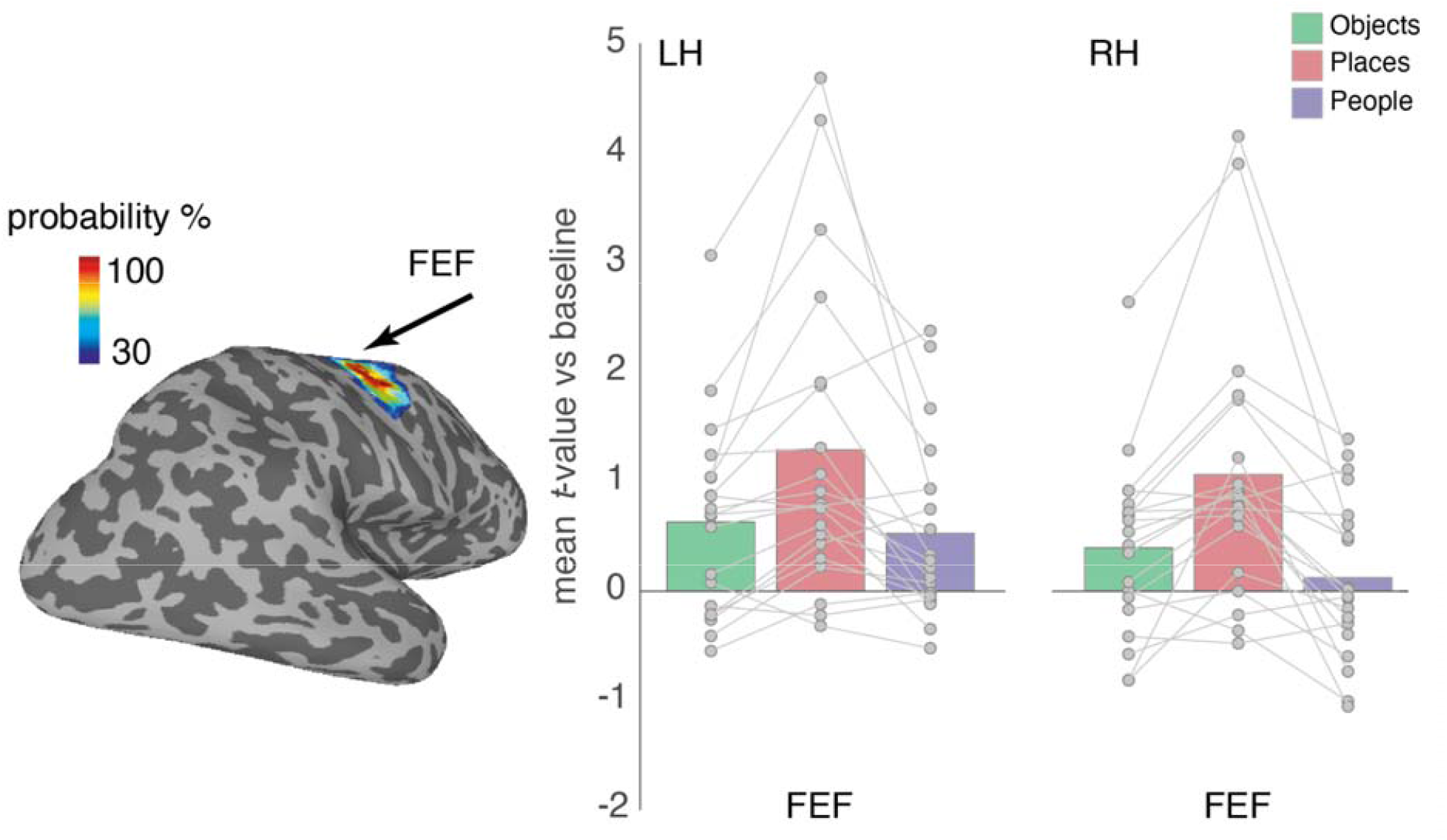
Place recall in the FEF. A lateral view of the right hemisphere is shown with the probabilistic mask for the FEF taken from (Wang et al., 2015) thresholded at 30% of the maximum probability. Bars represent the mean response to all categories from this FEF ROI in both hemispheres. Individual participant data points are plotted and connected. Place recall maximally drove responses in both hemispheres. The responses during both people and object recall were weaker but largely equivalent.

### Object recall effects in posterior parietal cortex

We also found significant category effects in posterior parietal cortex, in and around the post-central sulcus, that were particularly prominent for the contrast of people versus object recall **(Figure 7)**. This region is posterior to primary motor (M1) and somatosensory (S1) cortices as well as BA 1 and 2, and superior to secondary somatosensory cortex. It does not seem to correspond directly with any previously identified regions. However, responses in this area have been associated with grasping, but not reaching (S. H. Frey et al., 2005; Konen et al., 2013) or touching (Cavina-Pratesi et al., 2010).

**Figure 7:**
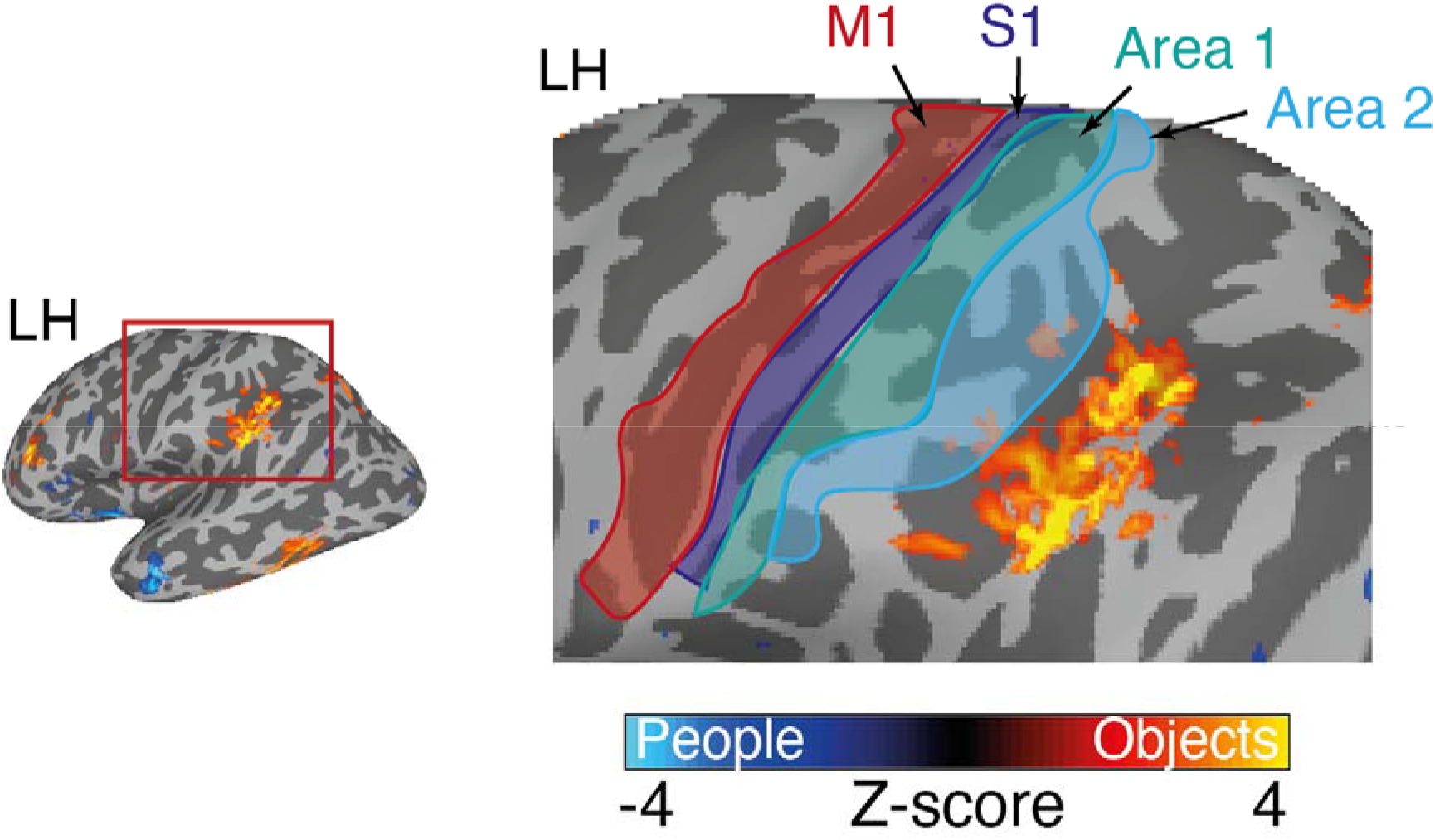
Object recall in posterior parietal cortex. A lateral view of the left hemisphere is shown with the posterior parietal cortex highlighted. The contrast of people versus objects is overlaid and enlarged to the right. This object recall is in the vicinity of the supramarginal gyrus, but is posterior of primary motor cortex (M1, red-outline), primary somatosensory cortex (S1, blue-outline), Area 1 (green-outline) and Area 2 (cyan-outline) taken from a publicly available parcellation (Glasser et al., 2016).

### Recall effects in early visual cortex

The contrast of people versus places (Figure 2) and our whole-brain analysis also hinted at recall effects within early visual cortex (V1-V3). Given the links between foveal/peripheral vision and face/scene processing we explored potential recall effects in foveal/peripheral EVC using population receptive field (pRF) data from an independent group of participants **(Figure 8A)**. Accordingly, these data were submitted to a three-way rmANOVA with Hemisphere, ROI (Foveal, Peripheral) and Category as within-participant factors. Only the main effect of Category (F(38)=38.35, p=0.0001) and the ROI x Category interaction (F(38)=13.14, p=0.00004) were significant (p>0.05, in all other cases). Given the non-significant effect of Hemisphere, these data were collapsed across hemisphere and resubmitted to a two-way rmANOVA with Category and ROI as factors. The main effect of ROI (F(19)=0.89, p=0.35) was not significant, but both the main effect of Category (F(38)=10.75, p=0.0001) and the ROI x Category interaction (F(38)=13.18, p=0.00004) were, reflecting on average larger responses within the peripheral ROI and positive responses during place recall but negative responses during both people and object recall. A series of paired t-tests (Bonferroni corrected, alpha=0.0083) revealed no differences between conditions in the foveal ROI (p>0.07, in all cases), but in the peripheral ROI responses during place recall were greater than recall of either people (t(19)=3.70, p=0.001) or objects (t(19)=4.41, p=0.0002). There were no differences between the recall of objects and people (t(19)=1.72, p=0.10).

**Figure 8:**
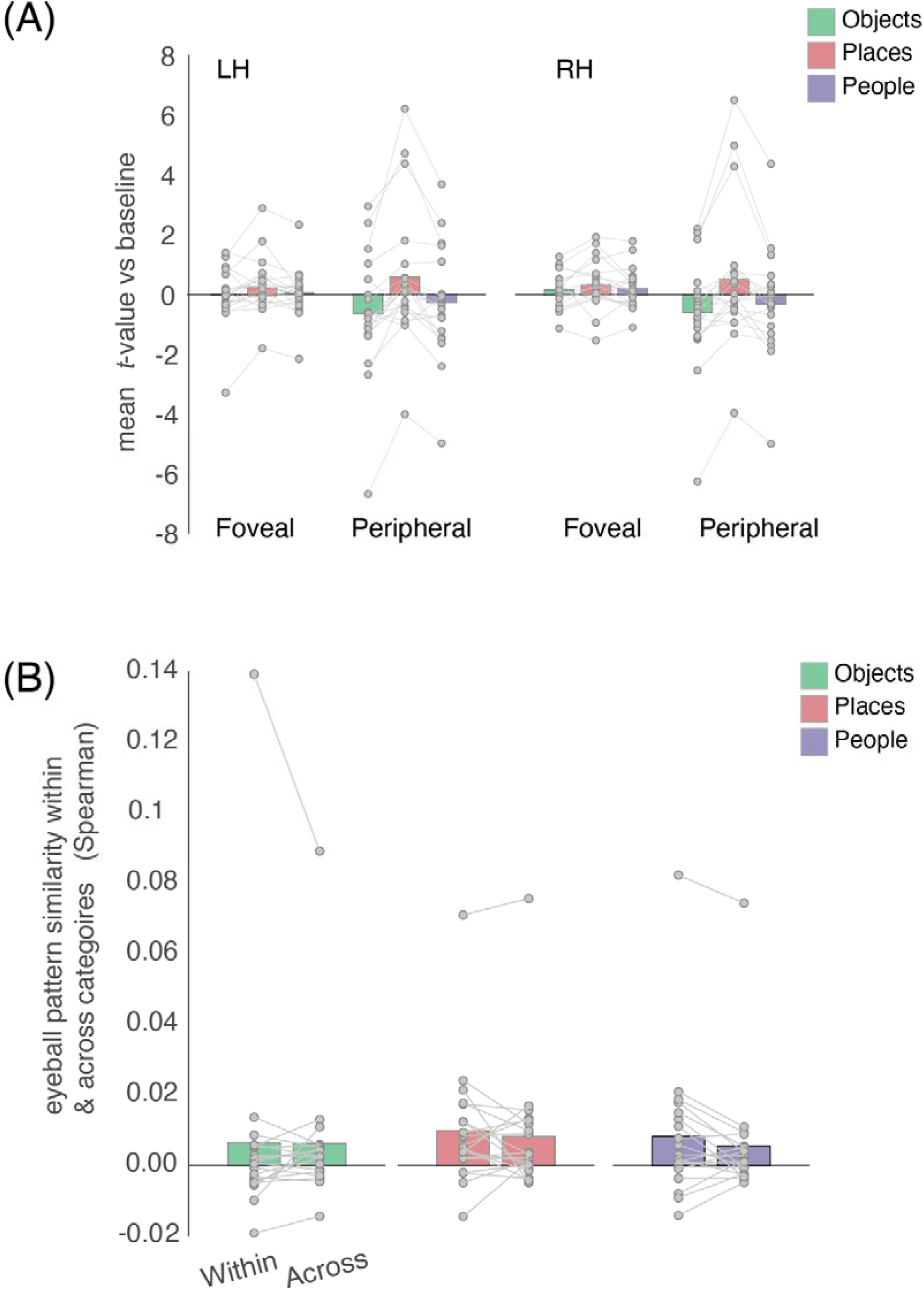
Recall effects in EVC & eyeball pattern similarity within & across categories. **(A)** Bars represent the mean response to all categories from foveal and peripheral EVC in both hemispheres. Overall, responses are largest during place recall in both foveal and peripheral EVC, with larger responses collectively in peripheral EVC. Individual participant data points are plotted and connected. **(B)** Bars represent the mean pattern similarity (Spearman) taken from the eyeballs both within and across categories. Individual participant data points are plotted and connected. In each case, there were no significant differences between the pattern similarity within and across categories.

### Eyeball-voxel analysis: an effect of category on eye movements?

Our exploratory whole-brain analysis revealed place recall-related activity in the FEF, a retinotopically organized region often associated with oculomotor control (Coiner et al., 2019; Mackey et al., 2017). We thus wondered if the observed FEF activity reflected eye movements that participants may have performed during recall. Indeed, prior work suggests that eye movements play a functional role in memory recall (Ryan & Shen, 2020). We reasoned that such recall-related eye movements may be more pronounced for places (or scenes) compared to objects or people, because of the often-complex arrangement and large number of features associated with a given place.

While we did not acquire eye-tracking data during scanning, recent work showed that eye movements affect the multi-voxel-pattern (MVP) of the eyeballs, and that the so inferable eye movements explain activity in the FEF (Frey & Nau et al., 2021). Therefore, to test if category-specific patterns in eye movements could explain our FEF results, we tested if the eyeball MVP in our data was indicative of the recalled category. We first extracted the eyeball MVP at each time point following established approaches (Frey & Nau et al., 2021), and averaged the MVP across functional volumes within each trial. We then computed the pairwise pattern dissimilarity between all trials of each category (within-category) as well as between trials of different categories (across-category). If participants moved their eyes more during recall of places compared to objects and people, we expected to find a weaker pattern dissimilarity for the within-category comparisons vs. the across-category comparisons. This was not the case **(Figure 8B)** (paired *t*-test of within versus across, *p* > 0.13, for all categories). However, each trial was 10 s long and the effect of eye movements on the MVP may have been lost when averaging across TRs. We therefore repeated the above analysis for each individual functional volume within each trial, again not finding an effect of category. Thus, this indirect measure of eye movements did not show a difference in viewing behavior between categories.

## Discussion

The current study was designed to accomplish two goals. The first was to provide an independent test of the people and place recall results reported previously (Silson, Steel, et al., 2019). The second was to test whether object-recall recruits distinct region(s) of MPC, akin to how perception of objects recruits distinct regions of VTC (i.e. pFS). Replicating our prior work to a striking degree, we identified an alternating pattern of people and place recall throughout MPC. However, despite strong predictions that were based on the similarities between VTC and MPC organization, object-recall effects were not observed within MPC itself. They were however observed in regions posterior and inferior to primary somatosensory cortex, particularly in the left hemisphere. Additional recall-effects were evident within the right pSTS during people recall and the FEF and EVC during place recall.

### Memory recall effects within MPC

Contrasting the responses elicited during people and place recall recapitulated original findings with a striking level of consistency. Such consistency was present not only within the topography and areal extent of the whole-brain activations, but also the pattern of responses within our MPC ROIs. The identification of four regions within MPC differentially recruited during people and place recall strongly supports our original report (Silson, Steel, et al., 2019) and recent independent fMRI (Deen & Freiwald, 2021) and neurophysiological evidence (Woolnough et al., 2020).

Within MPC itself we found no evidence for distinct recruitment during object recall either at the whole-brain or ROI analysis levels, despite strong predictions based on the otherwise parallel functional organization of VTC. Naturally, the absence of an effect is difficult to interpret, and it does not rule out that under other experimental settings MPC may show object selectivity during recall. In the following, we discuss several factors that could have contributed to this null result. First, recalling personally familiar people (e.g., Mother) and places (e.g., Childhood home) conceivably evokes stronger semantic, contextual, and emotional associations than the recall of familiar objects (e.g., Wallet). It is possible that the strong people and place responses and lack of object-recall within MPC reflect these differences in emotional/contextual and/or semantic importance (Bar & Aminoff, 2003). Unfortunately, we did not collect subjective ratings of the emotional/semantic significance of recalled items in order to test this possibility directly but taking these and other factors into account is a key goal of our future work. Second, it is possible that our internal representations of objects, particularly the types of objects selected by our participants (e.g., Watch, Wallet) are evolutionarily too recent relative to representations of people and places to recruit distinct regions within MPC. Unlike representations of people and places, our species’ experience with these objects comes solely from the last few hundred years (if that). Third, previous work put forward an organizational framework for understanding MPC that reflected the medial-lateral axis of VTC (Silson, Steel, et al., 2019). Although several different dimensions are thought to be represented along this axis, including eccentricity (Levy et al., 2001; Hasson et al., 2002), animacy (Konkle & Caramazza, 2013), and real-world size (Konkle & Oliva, 2012), categorical preference is arguably the strongest and most robust organization, being present even in infancy (Deen et al., 2017; Kosakowski et al., 2021). Within this organization, responses to objects are generally weaker than those elicited by either faces or scenes. Taken from the perspective of a functional link between VTC and MPC, it is possible that the absence of object-recall effects within MPC reflects this reduced responsiveness in VTC relative to people and places. Along these lines, it is possible that MPC represents the most salient distinctions within VTC only, which might facilitate the accurate recollection of information during memory. It is well known that contrasting faces and scenes produces an antagonistic response within FFA and PPA, respectively. Indeed, across the commonly tested visual object categories, faces reliably produce the largest positive response in FFA but simultaneously the weakest response in PPA and vice-versa. Such a salient distinction in VTC could therefore be reflected in the prominence of people and place recall reported here and previously within MPC (Silson, Steel, et al., 2019; Woolnough et al., 2020; Steel et al., 2021; Bainbridge et al., 2021; Deen & Freiwald, 2021).

### FEF, place-recall & the role of eye-movements

Interestingly, we observed stronger recall activity in the FEF for places compared to people or objects, which we reasoned may reflect a difference in viewing behavior between these categories. This would be in line with the proposed role of the FEF in oculomotor control (Mackey et al., 2017; Coiner et al., 2019), and with the notion that eye movements support memory recall (Ryan & Shen, 2020). While our imaging results predicted that participants performed more eye movements when recalling places compared to the other categories, we did not have concurrent eye tracking data to test this prediction directly. However, prior work showed that eye movements affect the voxel pattern of the eyeballs (Frey & Nau et al., 2021). We therefore used representational similarity analysis to test if the eyeball voxels carried information about category. While this was not the case in our data, we would like to emphasize that there are multiple potential reasons why. An obvious one is that there might not have been a difference in viewing behavior between categories. However, our task, data and analysis measures were also not optimal to find these effects even if they were present. First, the word cue remained on screen for the entire duration of the trial, meaning that one would expect a certain amount of eye movements in all trials irrespective of category. The effects we were looking for were therefore likely very small. Second, our repetition time of 2.5 s may have simply been too long for picking up these small effects, because eye movements occur at a much higher rate. Even if there was an effect of category, it may have averaged out on the level of the functional volumes. Given these possible explanations, we feel it would be wrong to conclude that viewing behavior does not contribute to our results, and the possible link between category specific recall, FEF activity and eye movements remains a key question for future work. For the present work, we conclude that we do not find evidence for an effect of eye movements on our neuroimaging results.

### People recall in right pSTS

Importantly, we identified people recall effects within pSTS in the right hemisphere, which we did not observe in our original report (Silson, Steel, et al., 2019). Using an independent group-based ROI for the face-selective pSTS, we found that responses were heavily negative during both place and object recall in pSTS whereas during people recall responses were close to zero on average. Importantly, the current study only used personally familiar stimuli during recall, whereas our prior work collapsed across both famous and personally familiar effects when contrasting people versus place recall. The previously reported advantage for recall of personally familiar over famous items potentially explains why the pSTS was identified in the current study and not our prior work. The fact that responses were less negative for the preferred category is reminiscent of the negative responses during visual perception of scenes and faces in MPC reported previously (Silson, Steel, et al., 2019).

### Object-recall effects

Interestingly, our whole-brain analyses highlighted category recall effects inferior and posterior to primary somatosensory cortex, that were particularly prominent when contrasting objects versus people. This object-recall response falls in the vicinity of the supramarginal gyrus and postcentral sulcus, which have been implicated in the representation of tools and grasping behavior (Johnson-Frey et al., 2005; McDowell et al., 2018). The majority of recalled objects in this study were small, graspable and manipulable. It is possible that recall of such items automatically generated associated motor representations, similar to what has been reported previously during the visual perception of familiar manipulable objects (Smith & Goodale, 2015). This interpretation is in-line with models that suggest the organization of category representations in the brain are grounded in action (and perceptual/emotional) systems (Martin, 2016).

### Recall effects in early visual cortex

At first glance, the fact that recall of places recruited peripheral portions of EVC to a greater extent than the recall of either people or objects is consistent with the association of scenes and faces/objects with peripheral and foveal visual processing, respectively. However, the increased peripheral activity during place recall could also reflect the fact that on average the place cues tended to contain almost twice as many characters and thus extend more into the periphery. Future studies in which the target word was presented briefly and then removed during recall could explore this effect more definitively as well as be better placed to assess any categorical effects in eye-movements during recall.

## Conclusions

Categorical memory recall effects within MPC appear restricted to those representing either people or places. The prevalence of people and place representations within MPC is consistent not only across studies from our group (Silson, Steel, et al., 2019; Steel et al., 2021; Bainbridge et al., 2021), but also independent studies using different imaging methodologies (Woolnough et al., 2020; Afzalian & Rajimehr, 2021; Deen & Freiwald, 2021). This contrasts MPC to VTC, possibly because only the most salient categorical representations within VTC (i.e. faces and places) are recapitulated within MPC in the context of memory. While our data provide no evidence for object-recall effects within MPC itself, we did observe such effects within the vicinity of regions implicated in representing tools and grasping. In sum, the functional organization of MPC is not a direct mirror image of the one of VTC, despite clear parallels surrounding the recall of places and people, which we replicated.

## References

Afzalian, N., & Rajimehr, R. (2021). Spatially Adjacent Regions in Posterior Cingulate Cortex Represent Familiar Faces at Different Levels of Complexity. The Journal of Neuroscience, 41(47), 9807–9826. https://doi.org/10.1523/JNEUROSCI.1580-20.2021

Bainbridge, W. A., Hall, E. H., & Baker, C. I. (2021). Distinct Representational Structure and Localization for Visual Encoding and Recall during Visual Imagery. Cerebral Cortex, 31(4), 1898–1913. https://doi.org/10.1093/cercor/bhaa329

Baldassano, C., Beck, D. M., & Fei-Fei, L. (2013). Differential connectivity within the Parahippocampal Place Area. NeuroImage, 75, 228–237. https://doi.org/10.1016/j.neuroimage.2013.02.073

Bar, M., & Aminoff, E. (2003). Cortical Analysis of Visual Context. Neuron, 38(2), 347–358. https://doi.org/10.1016/S0896-6273(03)00167-3

Benoit, R. G., & Schacter, D. L. (2015). Specifying the core network supporting episodic simulation and episodic memory by activation likelihood estimation. Neuropsychologia, 75, 450–457. https://doi.org/10.1016/j.neuropsychologia.2015.06.034

Cavina-Pratesi, C., Monaco, S., Fattori, P., Galletti, C., McAdam, T. D., Quinlan, D. J., Goodale, M. A., & Culham, J. C. (2010). Functional Magnetic Resonance Imaging Reveals the Neural Substrates of Arm Transport and Grip Formation in Reach-to-Grasp Actions in Humans. Journal of Neuroscience, 30(31), 10306–10323. https://doi.org/10.1523/JNEUROSCI.2023-10.2010

Chrastil, E. R. (2018). Heterogeneity in human retrosplenial cortex: A review of function and connectivity. Behavioral Neuroscience, 132(5), 317–338. https://doi.org/10.1037/bne0000261

Coiner, B., Pan, H., Bennett, M. L., Bodien, Y. G., Iyer, S., O’Neil-Pirozzi, T. M., Leung, L., Giacino, J. T., & Stern, E. (2019). Functional neuroanatomy of the human eye movement network: A review and atlas. Brain Structure and Function, 224(8), 2603–2617. https://doi.org/10.1007/s00429-019-01932-7

Cox, R. W. (1996). AFNI: Software for Analysis and Visualization of Functional Magnetic Resonance Neuroimages. Computers and Biomedical Research, 29(3), 162–173. https://doi.org/10.1006/cbmr.1996.0014

Deen, B., & Freiwald, W. A. (2021). Parallel systems for social and spatial reasoning within the cortical apex [Preprint]. Neuroscience. https://doi.org/10.1101/2021.09.23.461550

Deen, B., Richardson, H., Dilks, D. D., Takahashi, A., Keil, B., Wald, L. L., Kanwisher, N., & Saxe, R. (2017). Organization of high-level visual cortex in human infants. Nature Communications, 8(1), 13995. https://doi.org/10.1038/ncomms13995

Epstein, R. A. (2008). Parahippocampal and retrosplenial contributions to human spatial navigation. Trends in Cognitive Sciences, 12(10), 388–396. https://doi.org/10.1016/j.tics.2008.07.004

Epstein, R. A., & Baker, C. I. (2019). Scene Perception in the Human Brain. Annual Review of Vision Science, 5(1), 373–397. https://doi.org/10.1146/annurev-vision-091718-014809

Favila, S. E., Kuhl, B. A., & Winawer, J. (2019). Perception and memory have distinct spatial tuning properties in human visual cortex [Preprint]. Neuroscience. https://doi.org/10.1101/811331

Frey, M., Nau, M., & Doeller, C. F. (2021). Magnetic resonance-based eye tracking using deep neural networks. Nature Neuroscience. https://doi.org/10.1038/s41593-021-00947-w

Frey, S. H., Vinton, D., Norlund, R., & Grafton, S. T. (2005). Cortical topography of human anterior intraparietal cortex active during visually guided grasping. Cognitive Brain Research, 23(2–3), 397–405. https://doi.org/10.1016/j.cogbrainres.2004.11.010

Gilmore, A. W., Nelson, S. M., Chen, H.-Y., & McDermott, K. B. (2018). Task-related and resting-state fMRI identify distinct networks that preferentially support remembering the past and imagining the future. Neuropsychologia, 110, 180–189. https://doi.org/10.1016/j.neuropsychologia.2017.06.016

Gilmore, A. W., Nelson, S. M., & McDermott, K. B. (2015). A parietal memory network revealed by multiple MRI methods. Trends in Cognitive Sciences, 19(9), 534–543. https://doi.org/10.1016/j.tics.2015.07.004

Glasser, M. F., Coalson, T. S., Robinson, E. C., Hacker, C. D., Harwell, J., Yacoub, E., Ugurbil, K., Andersson, J., Beckmann, C. F., Jenkinson, M., Smith, S. M., & Van Essen, D. C. (2016). A multi-modal parcellation of human cerebral cortex. Nature, 536(7615), 171–178. https://doi.org/10.1038/nature18933

Hassabis, D., Kumaran, D., & Maguire, E. A. (2007). Using Imagination to Understand the Neural Basis of Episodic Memory. Journal of Neuroscience, 27(52), 14365–14374. https://doi.org/10.1523/JNEUROSCI.4549-07.2007

Hasson, U., Levy, I., Behrmann, M., Hendler, T., & Malach, R. (2002). Eccentricity Bias as an Organizing Principle for Human High-Order Object Areas. Neuron, 34(3), 479–490. https://doi.org/10.1016/S0896-6273(02)00662-1

Johnson-Frey, S. H., Newman-Norlund, R., & Grafton, S. T. (2005). A Distributed Left Hemisphere Network Active During Planning of Everyday Tool Use Skills. Cerebral Cortex, 15(6), 681–695. https://doi.org/10.1093/cercor/bhh169

Kanwisher, N., & Dilks, D. D. (2013). The functional organization of the ventral visual pathway in humans. In The new visual neurosciences.

Kim, H. (2013). Differential neural activity in the recognition of old versus new events: An Activation Likelihood Estimation Meta-Analysis. Human Brain Mapping, 34(4), 814–836. https://doi.org/10.1002/hbm.21474

Konen, C. S., Mruczek, R. E. B., Montoya, J. L., & Kastner, S. (2013). Functional organization of human posterior parietal cortex: Grasping- and reaching-related activations relative to topographically organized cortex. Journal of Neurophysiology, 109(12), 2897–2908. https://doi.org/10.1152/jn.00657.2012

Konkle, T., & Caramazza, A. (2013). Tripartite Organization of the Ventral Stream by Animacy and Object Size. Journal of Neuroscience, 33(25), 10235–10242. https://doi.org/10.1523/JNEUROSCI.0983-13.2013

Konkle, T., & Oliva, A. (2012). A Real-World Size Organization of Object Responses in Occipitotemporal Cortex. Neuron, 74(6), 1114–1124. https://doi.org/10.1016/j.neuron.2012.04.036

Kosakowski, H. L., Cohen, M. A., Takahashi, A., Keil, B., Kanwisher, N., & Saxe, R. (2021). Selective responses to faces, scenes, and bodies in the ventral visual pathway of infants. Current Biology, S0960982221015086. https://doi.org/10.1016/j.cub.2021.10.064

Kundu, P., Brenowitz, N. D., Voon, V., Worbe, Y., Vertes, P. E., Inati, S. J., Saad, Z. S., Bandettini, P. A., & Bullmore, E. T. (2013). Integrated strategy for improving functional connectivity mapping using multiecho fMRI. Proceedings of the National Academy of Sciences, 110(40), 16187–16192. https://doi.org/10.1073/pnas.1301725110

Kundu, P., Inati, S. J., Evans, J. W., Luh, W.-M., & Bandettini, P. A. (2012). Differentiating BOLD and non-BOLD signals in fMRI time series using multi-echo EPI. NeuroImage, 60(3), 1759–1770. https://doi.org/10.1016/j.neuroimage.2011.12.028

Levy, I., Hasson, U., Avidan, G., Hendler, T., & Malach, R. (2001). Center–periphery organization of human object areas. Nature Neuroscience, 4(5), 7.

Mackey, W. E., Winawer, J., & Curtis, C. E. (2017). Visual field map clusters in human frontoparietal cortex. ELife, 6, e22974. https://doi.org/10.7554/eLife.22974

Martin, A. (2016). GRAPES—Grounding representations in action, perception, and emotion systems: How object properties and categories are represented in the human brain. Psychonomic Bulletin & Review, 23(4), 979–990. https://doi.org/10.3758/s13423-015-0842-3

McDowell, T., Holmes, N. P., Sunderland, A., & Schürmann, M. (2018). TMS over the supramarginal gyrus delays selection of appropriate grasp orientation during reaching and grasping tools for use. Cortex, 103, 117–129. https://doi.org/10.1016/j.cortex.2018.03.002

Peer, M., Salomon, R., Goldberg, I., Blanke, O., & Arzy, S. (2015). Brain system for mental orientation in space, time, and person. Proceedings of the National Academy of Sciences, 112(35), 11072–11077. https://doi.org/10.1073/pnas.1504242112

Ryan, J. D., & Shen, K. (2020). The eyes are a window into memory. Current Opinion in Behavioral Sciences, 32, 1–6. https://doi.org/10.1016/j.cobeha.2019.12.014

Saad, Z. S., & Reynolds, R. C. (2012). SUMA. NeuroImage, 62(2), 768–773. https://doi.org/10.1016/j.neuroimage.2011.09.016

Silson, E. H., Gilmore, A. W., Kalinowski, S. E., Steel, A., Kidder, A., Martin, A., & Baker, C. I. (2019). A posterior–anterior distinction between scene perception and scene construction in human medial parietal cortex. Journal of Neuroscience, 39(4), 705–717. https://doi.org/10.1523/JNEUROSCI.1219-18.2018

Silson, E. H., Steel, A., Kidder, A., Gilmore, A. W., & Baker, C. I. (2019). Distinct subdivisions of human medial parietal cortex support recollection of people and places. ELife, 8, e47391. https://doi.org/10.7554/eLife.47391

Smith, F. W., & Goodale, M. A. (2015). Decoding Visual Object Categories in Early Somatosensory Cortex. Cerebral Cortex, 25(4), 1020–1031. https://doi.org/10.1093/cercor/bht292

Steel, A., Billings, M. M., Silson, E. H., & Robertson, C. E. (2021). A network linking scene perception and spatial memory systems in posterior cerebral cortex. Nature Communications, 12(1), 2632. https://doi.org/10.1038/s41467-021-22848-z

Szpunar, K. K., Watson, J. M., & McDermott, K. B. (2007). Neural substrates of envisioning the future. Proceedings of the National Academy of Sciences, 104(2), 642–647. https://doi.org/10.1073/pnas.0610082104

Vilberg, K. L., & Rugg, M. D. (2008). Memory retrieval and the parietal cortex: A review of evidence from a dual-process perspective. Neuropsychologia, 46(7), 1787–1799. https://doi.org/10.1016/j.neuropsychologia.2008.01.004

Wagner, A. D., Shannon, B. J., Kahn, I., & Buckner, R. L. (2005). Parietal lobe contributions to episodic memory retrieval. Trends in Cognitive Sciences, 9(9), 445–453. https://doi.org/10.1016/j.tics.2005.07.001

Wang, L., Mruczek, R. E. B., Arcaro, M. J., & Kastner, S. (2015). Probabilistic Maps of Visual Topography in Human Cortex. Cerebral Cortex, 25(10), 3911–3931. https://doi.org/10.1093/cercor/bhu277

Weiner, K. S., Golarai, G., Caspers, J., Chuapoco, M. R., Mohlberg, H., Zilles, K., Amunts, K., & Grill-Spector, K. (2014). The mid-fusiform sulcus: A landmark identifying both cytoarchitectonic and functional divisions of human ventral temporal cortex. NeuroImage, 84, 453–465. https://doi.org/10.1016/j.neuroimage.2013.08.068

Woolnough, O., Rollo, P. S., Forseth, K. J., Kadipasaoglu, C. M., Ekstrom, A. D., & Tandon, N. (2020). Category Selectivity for Face and Scene Recognition in Human Medial Parietal Cortex. Current Biology, 30(14), 2707-2715.e3. https://doi.org/10.1016/j.cub.2020.05.018

